# The genome of New Zealand trevally (Carangidae: Pseudocaranx georgianus) uncovers a XY sex determination locus

**DOI:** 10.1101/2021.04.25.441282

**Authors:** Mike Ruigrok, Andrew Catanach, Deepa Bowatte, Marcus Davy, Roy Storey, Noémie Valenza-Troubat, Elena López-Girona, Elena Hilario, Matthew J. Wylie, David Chagné, Maren Wellenreuther

## Abstract

**Background:** The genetic control of sex determinism in teleost species is poorly understood. This is partly because of the diversity of sex determining mechanisms in this large group, including constitutive genes linked to sex chromosomes, polygenic constitutive mechanisms, environmental factors, hermaphroditism, and unisexuality. Here we use a *de novo* genome assembly of New Zealand silver trevally (*Pseudocaranx georgianus*) together with whole genome sequencing to detect sexually divergent regions, identify candidate genes and develop molecular makers.

**Results:** The *de novo* assembly of an unsexed trevally (Trevally_v1) resulted in an assembly of 579.4 Mb in length, with a N50 of 25.2 Mb. Of the assembled scaffolds, 24 were of chromosome scale, ranging from 11 to 31 Mb. A total of 28416 genes were annotated after 12.8% of the assembly was masked with repetitive elements. Whole genome re-sequencing of 13 sexed trevally (7 males, 6 females) identified sexually divergent regions located on two scaffolds, including a 6 kb region at the proximal end of chromosome 21. Blast analyses revealed similarity between one region and the aromatase genes *cyp19 (a1a/b)*. Males contained higher numbers of heterozygous variants in both regions, while females showed regions of very low read-depth, indicative of deletions. Molecular markers tested on 96 histologically-sexed fish (42 males, 54 females). Three markers amplified in absolute correspondence with sex.

**Conclusions:** The higher number of heterozygous variants in males combined with deletions in females support a XY sex-determination model, indicating the trevally_v1 genome assembly was based on a male. This sex system contrasts with the ZW-type sex system documented in closely related species. Our results indicate a likely sex-determining function of the *cyp19b*-like gene, suggesting the molecular pathway of sex determination is somewhat conserved in this family. Our genomic resources will facilitate future comparative genomics works in teleost species, and enable improved insights into the varied sex determination pathways in this group of vertebrates. The sex marker will be a valuable resource for aquaculture breeding programmes, and for determining sex ratios and sex-specific impacts in wild fisheries stocks of this species.

## Introduction

The genetic basis of sex determination (SD) in animals has long fascinated researchers due to the relationship of this trait with reproduction and Darwinian fitness (Mank et al. 2006; Mank and Avise 2009). Traditionally, sex determination was assumed to be a relatively conserved trait across vertebrates. However, recent research on teleost fishes has shown that this is not the case, and that teleosts display a remarkable diversity in the ways sex is determined. These different mechanisms, which include heterogamety for males (males XY females XX) or heterogamety for females (males ZZ females ZW), multiple sex chromosomes and genes determining sex, environmental influences (temperature-dependent), epigenetic sex determination and hermaphroditism, have each originated numerous and independent times in teleosts (Volff 2005; Mank and Avise 2009; Piferrer 2013). The evolutionary lability of SD, and the corresponding rapid rate of turn-over among different modes, makes the teleost clade an excellent model to test theories regarding the evolution of SD adaptations (Sandra and Norma 2010; Yamamoto et al. 2019).

Teleosts consist of over 30,000 species, making them the largest group of vertebrates (Nelson et al. 2016). This diversity in species corresponds to a high phenotypic diversity and associated capacity of adaptation in physiological, morphological and behavioural traits. Reproductive systems vary largely, and strategies range from gonochorism, proterandrous, protogynous and simultaneous hermaphroditism (Devlin and Nagahama 2002). These reproductive strategies emerged independently in different lineages demonstrating a polyphyletic origin. Looking across fish families and genera, the genetic basis of tSD can be profoundly different, and can also be determined entirely by external factors, e.g. social structure or attainment of a critical age (Pla et al. 2021). Importantly, it should be noted that for most fish species it is unknown how sex is genetically determined and what the genetic architecture is (e.g. monogenic vs polygenic architecture).

The New Zealand silver trevally *Pseudocaranx georgianus* (hereafter referred to as trevally) also known as ‘araara’, its indigenous Māori name, is a teleost fish species of the family Carangidae. This family consists of approximately 30 genera which together contain for around 151 species worldwide (Fricke et al. 2018), yet SD has only been studied in a few species of this family. These studies have revealed that all of the carangids species are gonochoristic and that SD is genetically controlled (Crabtree et al. 2002; Devlin and Nagahama 2002; Graham and Castellanos 2005), which means that each individual has to be either a genetic male or female and is incapable of changing sex. Trevally is a pelagic species and abundant in the coastal waters of Oceania, spanning from the coastal regions of the North Island and the top of the South Island of New Zealand to southern Australia (Kailola 1993; Smith-Vaniz and Jelks 2006; Papa et al. 2020). The fish grows to a maximum length of 1.2 m and 18 kg, and can reach 25 years (Bray 2020). Their bodies are elongated, with the upper portion being bluish-silver, the lower portion of the fish is silver and the sides are yellow silver in colour (see Figure 1A, Bray 2020). They commonly school with size similar individuals and forage on plankton and bottom invertebrates (Smith-Vaniz and Jelks 2006).

**Figure 1.**
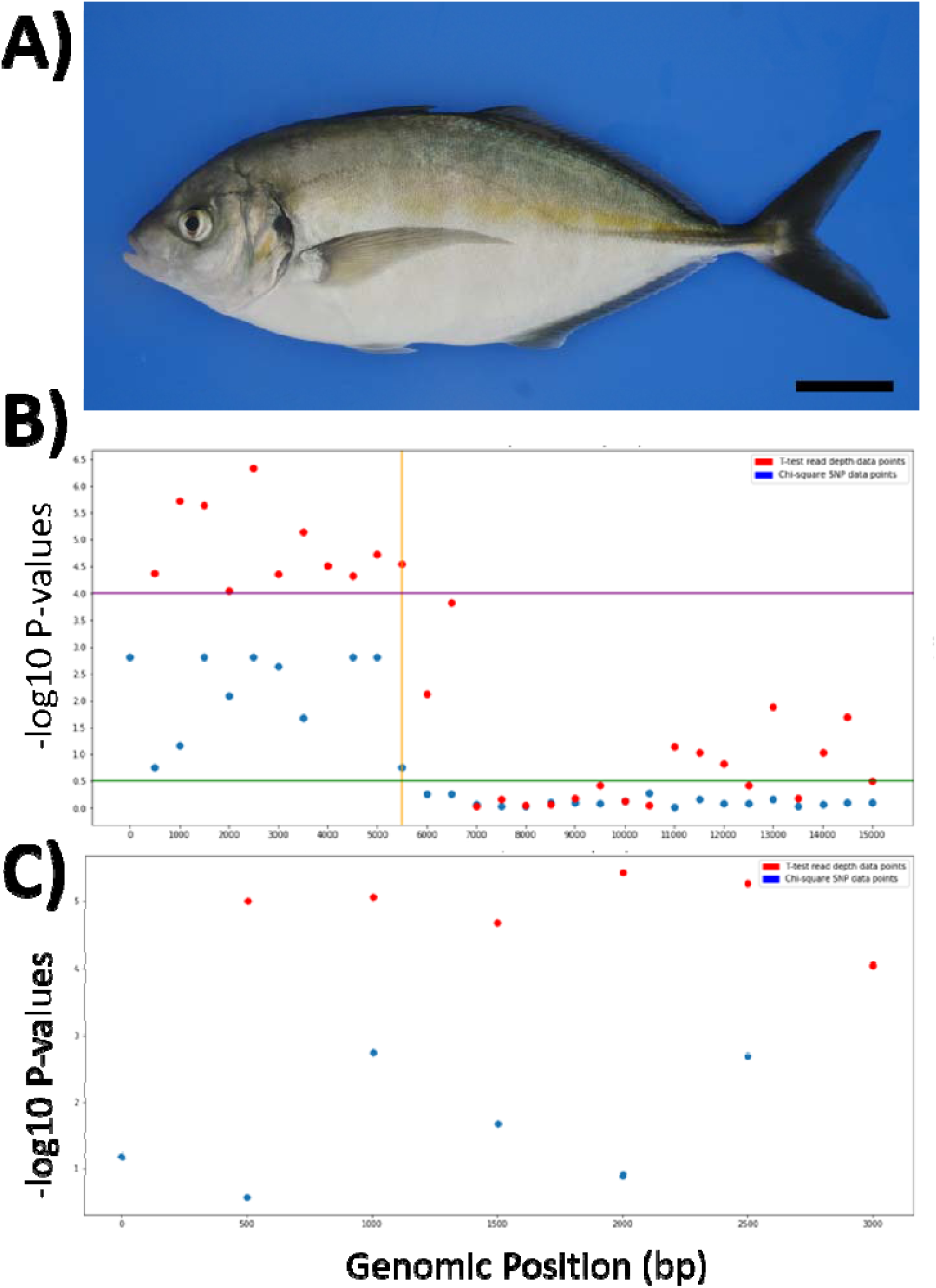
**Panel A)** shows a trevally individual from the breeding programme at Plant and Food Research in Nelson, New Zealand. The black scale bar is 5 cm. **Panel B)** shows the −log 10 P-values on the y-axis versus the genomic position (bp) on chromosome 21 (001800 scaffold) on the x axis in windows of 500 bases. **Panel C)** shows the −log 10 P-values on the y-axis versus the genomic position (bp) scaffold 000374 on the x axis in windows of 500 bases. −log 10 p-values were determined from p-values derived from t-tests (R) and from chi-square contingency tests (Python). A purple line is drawn at a −log 10 p-value of 4.0 for chromosome 21 to indicate a threshold for divergent windows for t-test read depth data points, a green line at a −log 10 p-value of 0.5 on chromosome 21 does the same for chi-square SNP data points. An orange line is drawn at a base position of 5500 to show where the SDR of chromosome 21 ends. Red dots indicate results from the t-test read depth analysis and blue dots SNP results from the chi-square test.

Trevally is a highly sought after sashimi species in Asia, and several countries are trying to establish aquaculture breeding programmes (e.g. Valenza-Troubat et al. in press). Adults of this species are sexually monomorphic externally, as observed in other carangids (Moriwake et al. 2001; Mylonas et al. 2004). Trevally have a firm musculature around their abdominal cavity, making manual sexing difficult. Thus, sex can typically only be determined subsequent to lethal sampling or by gonopore cannulation to retrieve a gonadal biopsy. This technique, however, can only be applied to broodstock in the advanced stages of gametogenesis shortly before or during the reproductive season and can injure the fish. Sexual maturation takes 3-4 years in captivity, meaning that sex information can only be gathered following that stage. Hence, understanding the genetic basis of SD in trevally would allow the design of molecular markers to facilitate sexing of the individuals early in life and in a less-invasive way.

The overarching goal of the study was to identify the genetic underpinnings of SD in trevally. To achieve this, we 1) *de novo* assembled a reference genome and 2) identified sexually-divergent genomic regions based on sequencing depth and variant detection using whole genome re-sequencing of male and female fish. Then, 3) candidate genes for SD were identified and 4) molecular markers were designed and validated using individuals sexed by gonadal histology. We discuss our findings about SD in this species and highlight the resulting applications, and compare them to other teleost species to draw general conclusions about SD in this group.

## Materials and Methods

### Broodstock collection and rearing of F_1_ offspring

The trevally samples used in this study were collected from a founding (F_0_) wild-caught captivity-acclimated population and a captive-bred (F_1_) generation produced by the New Zealand Institute for Plant and Food Research Limited (PFR) in Nelson, New Zealand. All fish were maintained under an ambient natural photoperiod and ambient water temperatures of filtered flow-through seawater. Fish were fed daily by hand to satiation on a diet consisting of a commercial pellet feeds (Skretting and/or Ridely) supplemented with frozen squid *(Nototodarus spp)* and an in-house mixed seafood diet enriched with vitamins.

F_0_ broodstock (n=21; mean body weight 1.4 ± 0.5 kg; mean fork length 44.3 ± 5.6 cm) were captured from two trawl fishing tows from the North Taranaki Bight (Lat. 3845267-Long. 17420626 and Lat. 3851887-Long. 17419780) during February 2012. Captured fish were transported to the Wakefield Key Finfish Facility (formerly operated by PFR in Nelson, New Zealand) and acclimated to a single 4400 L tank. Broodstock (remaining n=19) were later transferred to the Maitai Finfish Facility (currently operated by PFR in Nelson, New Zealand) in 2014 and were acclimated to a single 13000 L tank.

Spawning was induced to produce the F_1_ population in December 2015 when the ambient water temperature had reached 17.9 °C. In brief, the tank was fitted with an external passive egg collector and broodstock (mean body weight 3.5 ± 0.9 kg; mean fork length 51.4 ± 3.7 cm) received an intraperitoneal injection of human chorionic gonadotropin (Chorulon®) at a target dose of 600 IU/ kg bodyweight. Following injection, two individuals died at three and five days post-injection. The egg collector was checked daily (between 8 am and 9 am) for spawned eggs. Egg release was first observed at 48 h days post-injection, and at approximately 7-9 days post-injection, 50 g of buoyant eggs were collected for incubation over three consecutive days (fertilisation rates and egg production metrics were unreported). Eggs were transferred to individual 450 L upwelling conical incubators supplied with gentle aeration and ambient seawater (temp range: 19.4-21.7°C). At hatching, F_1_ larvae were combined into a single 5,000L tank where they were fed enriched rotifers *(Brachionus plicatilis)*, newly hatched artemia nauplii (*Artemia franciscana*) and reared using a semi-static green water rearing protocol. When larvae reached a notochord length of ∼7 mm, they were weaned onto fully enriched fully enriched *Artemia salina* followed by dry crumb (O.range, INVE Aquaculture) and an in-house wet-diet consisting of minced seafood. At approximately 77 days post-hatching, all fish were transferred into a single 63,000 L tank for on-growing under the husbandry conditions described above.

In 2017, a single two-year-old F_1_ juvenile was sampled for the genome assembly (section Genome sequencing and assembly), while 7 additional fish were sampled to quantify tissue specific RNA expression and to annotate the genome (tissues sampled: skin, white muscle, gill, liver, kidney, brain and heart tissues) (section RNA extraction for transcriptome sequencing). Three-year-old F_1_ individuals (n=96) were lethally sexed and sampled in 2018 and used for validation of the sex marker.

### Genome sequencing and assembly

#### DNA extraction for genome sequencing

Liver and heart tissues were collected from a two-year-old F_1_ individual and immediately preserved in RNAlater (Sigma-Aldrich, St Louis, USA) as recommended by the manufacturer. Total genomic DNA was extracted from a subsample of each white muscle tissue (∼20 mg) placed in 750 µL of CTAB buffer (2% hexadecyltrimethyl ammonium bromide, 2% polyvinyl pyrrolidinone K40, 2 M NaCl, 25 mM EDTA, 100 mM Tris-HCl pH 8.0) containing 2% β-mercaptoethanol. Tissue was disrupted by hand with a sterile plastic pestle until a homogeneous mixture was obtained. Homogenised samples were then extracted twice by vortexing with one volume of chloroform:isoamyl alcohol (CIA, 24:1) for 10 sec to remove the denatured proteins and centrifuged at 16,000 *g* for 5 min to separate solid and aqueous phases. The final aqueous phase was then transferred to a new 1.5 mL screw capped tube and the genomic DNA was precipitated by adding 0.7 volumes of room temperature isopropanol and left to precipitate at −20 °C for at least 1 h. DNA was collected by centrifugation at room temperature for 10 min at 16,000 *g*. The DNA pellet was washed with 1 mL of 70% ethanol (v/v). After all traces of ethanol were removed by air drying, DNA was slowly dissolved in 100 µL sterile TE buffer (10 mM Tris-HCl pH 7.5, 1 mM EDTA) at 4 °C overnight. RNA was removed by adding 4 µL of RNase A (100 mg/µL) to the DNA and incubating for 5 min at room temperature. Following RNase treatment, samples were subjected to a final CTAB extraction as described above. The final DNA pellet was dissolved in 100 µL of TE buffer and quantified by fluorescence (Qubit™ HS dsDNA kit Invitrogen). DNA quality was assessed by standard gel electrophoresis (1% agarose in 1X TAE buffer, 40 mM Tris-acetate, 1 mM EDTA at pH 8.3) and by pulse field gel electrophoresis in 1% Certified™ Megabase Agarose (Bio-Rad) in 1X TAE buffer. The average fragment size of the DNA was 40 kb

### Genome assembly

#### Short-insert library preparation, sequencing, and assembly

Dovetail Genomics (Scotts Valley, CA, USA) was contracted to conduct the de novo sequencing project, which consisted of a short insert library and two long range libraries (Hi-C and Chicago). The Illumina short insert library was prepared with randomly fragmented DNA according to the manufacturer’s instructions. The library was sequenced on an Illumina HiSeq X platform using paired-end (PE) 150 bp sequencing. The data were trimmed for low-quality bases and adapter contamination using Trimmomatic and Jellyfish (Marçais and Kingsford 2011) with an in-house software to profile the short insert reads at a variety of k-mer values (25, 43, 55, 85 and 109) to estimate the genome size, and fit negative binomial models to the data. The resulting profiles suggested a k-mer size of 43 was optimal for assembly. The contigs were assembled into scaffolds using Meraculous (Chapman et al. 2011), with a k-mer size of 43, a minimum k-mer frequency of 12, and the diploid nonredundant haplotigs mode.

#### Chicago library preparation and sequencing

Second, following the *de novo* assembly with Meraculous, a Chicago library was prepared according to the methods described in Putnam et al. (2016). Briefly, ∼ 500 ng of high molecular weight genomic DNA was reconstituted *in vitro* into chromatin and subsequently fixed with formaldehyde. The fixed chromatin was then digested with DpnII, the 5′ overhangs were filled in with biotinylated nucleotides and free blunt ends were ligated. After ligation, crosslinks were reversed and the DNA was purified from any protein. The purified DNA was them treated to remove biotin that was not internal to ligated fragments and the resulting DNA was sheared to ∼350 bp mean fragment size using a Bioruptor Pico. Sequencing libraries were prepared from the sheared DNA using NEBNext Ultra enzymes (New England Biolabs, Inc.) and Illumina-compatible adapters. The biotin-containing fragments were isolated using streptavidin beads before PCR enrichment of each library. The amplified libraries were finally sequenced on an Illumina HiSeq X platform using PE 150 reads to approximately 90X depth.

#### Dovetail Hi-C library preparation and sequencing (multiple libraries)

Third, a Dovetail Hi-C library was prepared from the heart tissue preserved in RNAlater following the procedures outlined in Lieberman-Aiden et al. (2009). Briefly, formaldehyde was used to fix chromatin in place in the nucleus, which was then extracted and digested with DpnII. The 5′ overhangs were filled with biotinylated nucleotides, and free blunt ends were ligated. After ligation, the crosslinks were reversed and the DNA was purified from remaining protein. Biotin that was not internal to ligated fragments was removed from the purified DNA, which was subsequently sheared to ∼350 bp mean fragment size using a Bioruptor Pico. The sequencing libraries were then prepared using NEBNext Ultra enzymes and Illumina-compatible adapters. Before PCR enrichment of the library, biotin-containing fragments were isolated using streptavidin beads. The resulting library was sequenced on an Illumina HiSeq X Platform using PE 150 reads to approximately 60X depth.

#### Assembly scaffolding with HiRise

To scaffold and improve the trevally *de novo* assembly, Dovetail staff input the Meraculous assembly, along with the shotgun reads, Chicago library reads, and Dovetail Hi-C library reads into the HiRise pipeline (Putnam et al. 2016) to conduct an iterative analysis. First, the shotgun and Chicago library sequences were aligned to the draft contig assembly using a modified SNAP read mapper (http://snap.cs.berkeley.edu). Second, the separations of Chicago read pairs mapped within draft scaffolds were analysed to produce a likelihood model for genomic distance between read pairs. This model was used to identify and break putative misjoins, score prospective joins, and make joins above a threshold. Finally, after aligning and scaffolding the draft assembly using the Chicago data, the Chicago assembly was aligned and scaffolded using Dovetail Hi-C library sequences following the same method. After scaffolding, the short-insert sequences were used to close remaining gaps between contigs where possible.

#### Assembly polishing and contiguity statistics

After receiving the assembly from Dovetail, *de novo* repeats were identified using RepeatModeler v1.0.11 (http://www.repeatmasker.org/RepeatModeler.html) with the NCBI search engine (rmblast version). Repeats were classified by RepeatModeler into simple, tandem and interspersed repeats and masked using RepeatMasker v4.0.5 (Smit and Hubley 2008).

### RNA extraction for transcriptome sequencing

Skin, white muscle, gill, liver, kidney, brain and heart tissues were collected from five randomly selected F_1_ and immediately placed in RNAlater. RNA was extracted from the five replicates with the CTAB buffer described above as follows: approximately 50 mg of tissue were processed as for the DNA preps until the aqueous phase was obtained after the second CIA extraction. At this point the aqueous phase was precipitated with 0.35 volumes of 8 M LiCl, mixed by inversion and incubated at 4 °C overnight. The RNA was collected by centrifugation at 16,000 *g* in a refrigerated micro centrifuge. The pellet was dissolved in 500 µL SSTE buffer (1 M NaCl, 0.5% SDS, 1 mM EDTA, 10 mm Tris-HCl pH 8.0) and extracted once with an equal volume of CIA. The aqueous phase was collected after 10 min centrifugation at 16,000 *g* and precipitated with 2 volumes of 100% ethanol. The RNA was collected by centrifugation at 16,000 *g* and washed with 1 mL of 70% ethanol, then air dried at room temperature for approximately 30 min. The RNA was dissolved in 200 µL sterile deionized water. The RNA quality was assessed by absorbance ratios (260/280 nm and 260/230 nm) and was quantified by absorbance at 260 nm. The samples were DNase-treated with the Ambion® TURBO DNA-free™ kit (Thermo-Fisher Scientific, Waltham, MS, USA) as directed by the manufacturer. The RNA quality and quantity were assessed by absorbance, as described above. The fragment-size distribution was assessed by capillary electrophoresis using a Fragment Analyzer (Advanced Analytical, Parkersburg, WV, USA), using the High Sensitivity RNA Analysis kit.

The RNA samples were sequenced at the Australian Genome Research Facility (AGRF). The best replicate from each tissue was selected for the transcriptome data set by preparing TrueSeq libraries (Illumina, San Diego, CA, USA). The rest of the samples were prepared with the Lexogen QuantSeq 3’mRNA kit (Lexogen, Wien, Austria) for the expression analysis data set.

### Genome annotation

Automated gene models were predicted using the BRAKER2 pipeline v2.1.0 (Hoff et al. 2018) with trevally RNA sequences and the trevally genome assembly as input. Gene and genome completeness were evaluated using BUSCO v3.0.2 (Simão et al. 2015) using the vertebrata_odb9 lineage set (containing 2586 genes). Functional annotations were assigned to the gene models using blastx (Aitschul 1990) to search for similarities between the translated transcriptome gene-locus models and a peptide database using 88,504 peptide sequences of *Danio rerio* and 39,513 peptide sequences of *Seriola lalandi* (downloaded from NCBI using E-utilities version 11.4, 7^th^ September 2020). The results from these searches were merged with species-specific genome-wide annotation for Zebrafish (*Danio rerio*) provided in the package org.Dr.eg.db (Carlson 2019), using Entrez stable gene identifiers (Maglott et al. 2007) and Genbank accessions to annotate BLASTX alignments of gene models. Common Gene Locus (gene model g1 .. g28000) from blast reports were also used to marry up Zebrafish, and Kingfish accession and description information.

### Whole genome sequencing of sexed F_0_ broodstock

Sampling of the 13 remaining broodstock (of the original 21) took place during February 2017. Fin tissue (fin clips) were placed directly into chilled 96% ethanol, heated to 80 °C for 5 minutes within 1 hour of collection, and then stored at −20 °C. Total genomic DNA was extracted as follows: approximately 20 mg of fin tissue were added to a mixture of 400 µL extraction buffer (0.4 M NaCl, 10 mM Tris-HCl pH 8.0, and 2 mM EDTA pH 8.0) and 80 µL of 10% SDS. The sample was incubated at 80°C for 5 min and immediately cooled on ice. Ten microliters of Proteinase K (10 mg/mL) were added and mixed by inversion. The sample was incubated at 56 °C for 1.5 h. The insoluble material was removed by centrifugation at 16,000 *g* for 5 min. The supernatant was transferred to a new micro centrifuge tube and the proteins were salted out by adding 320 µL 5 M NaCl and mixing by inversion. The denatured proteins were collected by centrifugation at 16,000 *g* for 5 min. The clear supernatant was transferred to a new micro centrifuge tube and the RNA was removed by adding 5 µL RNase A 100 µg/µL, incubated at room temperature for 5 min. The sample was centrifuged at 16,000 *g* for 5 min and the supernatant was transferred to a new micro centrifuge tube. The DNA was precipitated with 525 µL isopropanol and incubated at −20°C overnight, prior to collection by centrifugation at 16,000 *g* for 10 min. The pellet was washed with 1 mL of 70% ethanol, dried and resuspended in 100 µL TE buffer. DNA quality was assessed by absorbance ratios (260/280 nm and 260/230 nm) and the quantified by fluorescence (Qubit™ HS dsDNA kit, Invitrogen). The integrity of the DNA was assessed by capillary electrophoresis using the High Sensitivity Genomic DNA Analysis kit on the Fragment Analyzer (Illumina). Short insert (300 bp) libraries (Illumina) were prepared and sequenced by AGRF (PE reads, 125 b long).

#### Whole genome sequence read alignment and variant detection

FASTQ files of reads belonging to the 13 sexed F_0_ broodstock were quality filtered using Trimmomatic v0.36 (Bolger et al. 2014) with a sliding window size of 4, a quality cut-off of 15 and the minimum read length set at 50. Filtered FASTQ files were aligned to the reference genome Trevally_v1 using BWA-MEM v0.7.17 (Li 2013). Aligned BAM files of two sequencing lanes per individual fish were merged using Samtools v1.7 (Li et al. 2009). Read groups were added and duplicates were removed from merged BAM files using Picard Tools v2.18.7, and sorted and indexed using Samtools. Variant calling was done on the whole cohort of 13 fish using freebayes-parallel v1.1.0 (https://github.com/freebayes).

### Genome-wide detection of sex-linked variants

Two strategies were used for detecting sex-linked regions using the re-sequencing data from the 13 sexed broodstock (Supplementary Figure 1). A read-depth based approach was employed by exploiting the difference in sex chromosome ploidy between males and females. For this, alignments to scaffolds shorter than 3000 bp were excluded from bam files using an in-house BASH script with AWK (Supplementary Material, Read_depth_analysis.ipynb). Read-depth was calculated per base using Samtools v1.7 (Li et al. 2009). A variant density approach was employed, searching for differences in SNPs and insertion-deletion (indels) density between males and females. To identify sexually divergent regions, the VCF data were converted to genotypes using the R package VCFR (R version: v4.3.3, package-version: v1.12.0). At each variant position, frequencies of genotypes were calculated for males and females, factorized per base position with the Python v3.6.5 factorize module and prepared in an array, which consisted of three columns: male frequency, female frequency and genotype variants in the form of integers (this included indels). This array increased or decreased in size based on the number of variants that were present on a given base position. This array was used as input for the Python chi2_contingency module (https://docs.scipy.org/doc/scipy/reference/generated/scipy.stats.chi2_contingency.ht ml), from which p-values and their corresponding scaffold and location were extracted and stored in a variant divergence file.

With R v3.5.0 all base positions from the variant divergence file (file with indel and SNP divergence per scaffold and location) were converted into windows of 500 bases and the mean p-value over these windows was calculated, the same was done for depth with the created depth files (the files that store depth values per scaffold). The windows of depth were then submitted in a t-test, after separating the windows by sex, with p-values as output. A fixation index (Fst) was calculated with VCFtools v0.1.14 (Danecek et al. 2011) with an Fst window size of 500 base pairs to create windows of the same genomic regions as the other tests, with the initial VCF file as input.

The 500 base pair windows with their corresponding p-values (from the read depth t-test and chi-square genotype test) and Fst (from the VCF file) were filtered by sorting these values from high to low based on their differential value with a BASH AWK script (Supplementary Material, Analysis_and_primer_creation.ipynb). The top windows were investigated with Integrative Genomics Viewer IGV v2.3 (Robinson et al. 2011) for a quick overview of the genomic differences in terms of depth and SNPs, from these sexually divergent regions were noted (big differences in depth or SNPs). The sexually divergent regions found were then investigated on their exact numbers of indel and hetero and homozygosity to infer the sex system. To do this, the sexually divergent regions and random regions of the same size from the earlier created VCF file with Bcftools v1.9 (Narasimhan et al. 2016) were indexed and extracted into VCF files that only contained the data of these regions, after which the data was converted to a genotype format from which a generic Python script counted deletions and hetero/homozygous SNP data relative to sex, which was later converted to a ratio by dividing the allele male count with the allele female count.

### Identification of candidate genes related to sex determination

Teleost SD candidate genes were identified and compiled from publications from 1998 and onwards using the search terms “sex determination, *Pseudocaranx georgianus*, Carangidae, Perciformes, teleost, fish in combination with sex determination and sex genes” in Google Scholar (parsed from 01/09/2019 to 01/10/2019). Sequences of candidate genes were downloaded from NCBI and used to query the trevally reference genome Trevally_v1 using BLASTN v2.2.25 (Chen et al. 2015), filtering for E-values < 1e-10, and alignment lengths and bit scores greater than 99.

#### Sex phenotyping for marker development

For the development and validation of a molecular sex marker in trevally, gonadal and fin tissues were collected from three-year-old F_1_ individuals (n=96). In brief, fish were subjected to complete sedation and euthanasia by overdose in anaesthetic (> 50 ppm AQUI-S^®^; Aqui-S New Zealand Ltd, Lower Hutt, New Zealand) followed by cervical dislocation with a sharp knife.

A fragment of gonadal tissue was dissected and fixed in a solution of 4% formaldehyde-1% glutaraldehyde for at least 48 h at 4 °C. Fixed samples were then dehydrated through an ethanol series before being embedded in paraffin (Paraplast, Leica Biosystems Richmond Inc, Richmond IL, USA). Serial sections cut to a thickness of 5 μm were obtained using a microtome (Leica RM2125RT, Leica Microsystems Nussloch GmbH, Germany) and stained in Gill 2 hematoxylin (Thermo Scientific Kalamazoo, MI, USA) and counterstained with eosin. Histological sections were examined under a light compound microscope (Olympus BX50) for the presence of oocytes or spermatogonia and photographed with a digital camera (Nikon DS-Ri2) to confirm the sex of each individual.

### Sex marker development and validation

Fin clips were collected from the 96 individual F_1_ fish and placed directly into chilled 96% ethanol, heated to 80 °C for 5 minutes and then stored in a −20 °C freezer. Total genomic DNA was extracted as described above. Three types of genetic markers were developed in the sex-linked regions. PCR primers were designed using the Primer3 v4.1.0 web application. Y-specific markers were designed using male sequences where there is an absolute deletion for females, so that PCR only amplifies the Y allele. Gene-based primers were designed with default parameters using the trevally ortholog of *cyp19a1a* from *Seriola lalandi* (HQ449733.1). PCR primers for High Resolution Melting (HRM) were designed around the sexually divergent SNPs by flanking the SNPs with 100 bp on each side.

HRM markers were screened using PCR conditions and mix described in Guitton et al. (2012) using genomic DNA extracted from fin clips of these fish. Y-allele specific and candidate gene-based markers were screened as sequence-characterized amplified regions (SCAR) markers as described in Bus et al. (2008). PCR primers conditions were first tested on eight individual samples to verify PCR amplification and presence (in males) absence (in females) polymorphism, then screened on the population of 96 sexed fish.

## Results

### Genome sequencing and assembly

In total, 412,758,157 paired-end Illumina short reads were generated from an F_1_ unsexed trevally, of which 97.4% were retained after trimming. K-mer analysis (k=43) resulted in 0.71% of heterozygous SNPs and an estimated genome size of 646 Mb.

The total input sequencing data pre-assembly was approximately 121 Gb, which is equivalent to 187.3× coverage.

The whole genome assembly yielded 2,006 scaffolds greater than 1 kb, for a total assembly size of 579.4 Mb (89% of estimated genome) and a N50 (scaffold) of 25.2 Mb. Of this total assembly, 574.8 Mb (99.2% of the total assembly and 88.8% of the k-mer estimated genome size) were assembled into 24 chromosome-size scaffolds ranging from 11 Mb to 31 Mb in length and corresponding to the expected karyotype of trevally (Table 1). The remaining scaffolds (<0.8% of the total assembly) that could not be anchored to pseudo-chromosomes were smaller, ranging from 1 kb to 51.2 kb in size.

**Table 1:** The 24 anchored trevally chromosomes (following Valenza-Troubat et al. in preparation) and the corresponding scaffold names and their respective lengths (bp). Note, that the scaffold_001800 is located on Chromosome 21, making this the sex chromosome.

| Anchored chromosome | Scaffold name | Length (bp) |
| --- | --- | --- |
| Chr01 | scaffold_000114 | 24151137 |
| Chr02 | scaffold_000147 | 28820665 |
| Chr03 | scaffold_001154 | 27945907 |
| Chr04 | scaffold_001701 | 22194978 |
| Chr05 | scaffold_000836 | 22952508 |
| Chr06 | scaffold_001460 | 31062029 |
| Chr07 | scaffold_000312 | 24419205 |
| Chr08 | scaffold_000453 | 26980729 |
| Chr09 | scaffold_001891 | 28251306 |
| Chr10 | scaffold_001025 | 26936208 |
| Chr11 | scaffold_001200 | 18865379 |
| Chr12 | scaffold_000118 | 26912610 |
| Chr13 | scaffold_001584 | 22008664 |
| Chr14 | scaffold_000429 | 25278635 |
| Chr15 | scaffold_001123 | 23296617 |
| Chr16 | scaffold_000874 | 26415528 |
| Chr17 | scaffold_000314 | 22912052 |
| Chr18 | scaffold_001965 | 19902634 |
| Chr19 | scaffold_001072 | 19893011 |
| Chr20 | scaffold_000070 | 24159416 |
| Chr21* | scaffold_001800 | 27791086 |
| Chr22 | scaffold_000657 | 25934954 |
| Chr23 | scaffold_000099 | 16759215 |
| Chr24 | scaffold_000328 | 11005663 |
| Total anchored |  | 574850136 |
| Unanchored scaffolds |  | 4556253 |
| Total anchored + unanchored |  | 579406389 |

\*: sex-linked

#### Repeat and gene annotation

A total of 12.8% of the genome was masked for repeats. BUSCO analysis of the anchored Trevally_v1 genome yielded a complete BUSCO score of 92.4% with 2364% being single copy and 27% being duplicated copies (134 were fragmented and 61 missing). In total, 28416 protein-coding gene models were detected.

#### Whole genome re-sequencing and sex-determining regions mapping

A total of 107.8 Gb of Illumina short reads were produced for the 13 trevally F0 broodstock individuals. In total 16,576890 variants were detected, including 14,355149 and 2,221741 SNPs and indels, respectively. Chi-square and t-tests both found significant SNPs, indels and depth differences on two scaffolds (Figure 2B and C). The chi-square test using SNPs and indels detected significant hits with a −log10P value of 0.58 to 2.73 for the 1 to 3,000 bp region of scaffold_000374 including 66 significant SNPs. Scaffold_000374 is a small unanchored scaffold of 3.7 kb in length. A −log10P value of 0.75 to 2.82 was found for the 1 to 6,000 bp region of scaffold_001800 (chromosome 21) including 78 SNPs. Scaffold_001800 is a large pseudo-chromosome scaffold of 27.7 Mb in length (denoting chromosome 21) and the sex-linked region is at the end of this scaffold. The t-test using read depth had a differential −log10P ranging from 4.04 to 5.42 across the 1 to 3,000 bp region of scaffold_000374 and a differential −log10P ranging from 4.05 to 6.33 for the 1 to 6,000 bp region of scaffold_001800 (chromosome 21).

**Figure 2.**
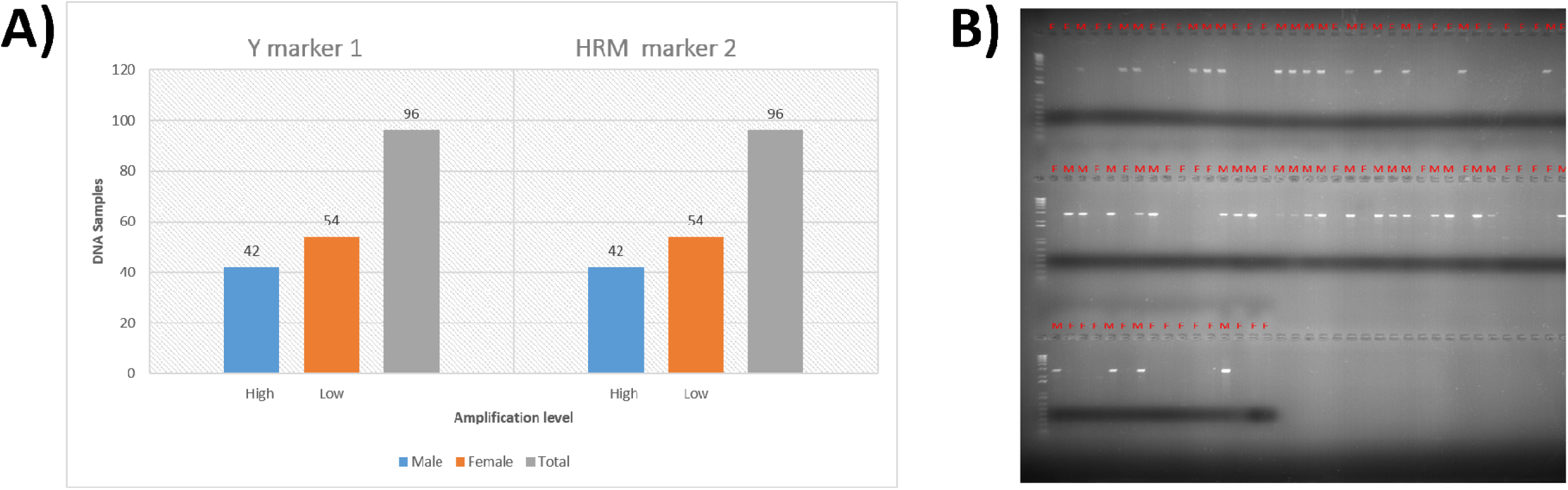
**Panel A)** Bar plot made from Y marker 1 and HRM marker 2 measured cycle amplification levels from running HRM on a LightCyler480(Roche) real time PCR instrument for males (blue) and females (orange). The Y-axis is the amount of DNA samples and the labels high or low indicate amplification levels. **Panel B**) 0.9% agarose gel stained with RedSafe^TM^, which represents PCR products (∼2.5 kb) from the sex-specific gene-wide marker. With red text above each slot containing either an M (Male) or F (Female).

#### Model for sex determination

The ratio of heterozygous and homozygous SNPs between males and females was 0.64 and 0.69 for scaffold_000374 and chromosome 21 (scaffold_001800), respectively (Table 2). This ratio is lower in non-linked (randomly selected and same sized) scaffolds (average of 0.05 and 0.11 respectively). There are more heterozygous variants in males than females in both candidate sex-linked regions (299 and 263 in favour of males), which is indicative of a XY system. Female sex-linked regions were identified by the presence of deletion variants (185 and 233 per *de novo* reference scaffold, respectively) on the female re-sequenced reads that were not called (or present) on male-reads. We found variant sites that were heterozygous for all the males and homozygous for all females (3 sites against scaffold_000374 and 8 on chromosome 21 (scaffold_001800)). In contrast, we did not find homozygote sites for the males that were heterozygous for the females (indicative of a XX/XY system), whilst the other way around 0 instances were detected (signs of a ZZ/ZW system).

**Table 2.** Overview of female and male differences in homo/heterozygosity and Indels. The blue highlighted cells contain data of the 000374 SDR region and the orange highlighted cells contain data of the 001800 (chromosome 21) SDR region. Next to the SDR data the highlighted regions also contain 3 randomly selected genomic regions of the same size and their data to show how differential the SDR is compared to a regular genomic region.

| Scaffold | Female DELS | Male DELS | Full hetero male<br>and homo female | Full homo female<br>and hetero male | Homo males | Homo females | Hetero males | Hetero females | Hetero<br>ratio(male/female) | Homo<br>ratio(male/female) | Ratio difference |
| --- | --- | --- | --- | --- | --- | --- | --- | --- | --- | --- | --- |
| Scaffold 001800 | 185 | 0 | 0 | 8 | 326 | 252 | 546 | 283 | 1.929328622 | 1.293650794 | 0.635677828 |
| Scaffold R1_6000 | 5 | 24 | 0 | 0 | 478 | 462 | 725 | 637 | 1.138147567 | 1.034632035 | 0.103515532 |
| Scaffold R2_6000 | 3 | 1 | 0 | 0 | 233 | 186 | 286 | 243 | 1.176954733 | 1.252688172 | 0.07573344 |
| Scaffold R3_6000 | 0 | 0 | 0 | 0 | 270 | 202 | 322 | 276 | 1.166666667 | 1.336633663 | 0.169966997 |
| Scaffold 000374 | 233 | 0 | 0 | 3 | 331 | 154 | 462 | 163 | 2.834355828 | 2.149350649 | 0.685005179 |
| Scaffold R1_3000 | 6 | 4 | 0 | 0 | 299 | 232 | 360 | 306 | 1.176470588 | 1.288793103 | 0.112322515 |
| Scaffold R2_3000 | 15 | 17 | 0 | 0 | 863 | 732 | 1180 | 1011 | 1.167161227 | 1.178961749 | 0.011800522 |
| Scaffold R3_3000 | 2 | 0 | 0 | 0 | 820 | 724 | 1015 | 868 | 1.169354839 | 1.132596685 | 0.036758154 |

### Identification of candidate genes related to sex determination

A total of 32 research publications were found and from these, a total of 132 candidate SD genes collated, of which 64 were unique (Supplementary Table 1). We used these 64 candidate genes as queries to search the trevally reference genome using BLAST and we produced 6119 matches (prior to filtering), of which 31 were found in the two sex-linked scaffolds with E-values ranging from 1.00E-25 to 9.00E-22 and high identity rates ranging from 78.98 to 88.07 (Table 3). Every hit within the sex-linked region impacted one of two genes; *Cyp19b* and *Cyp19a1a*.

**Table 3.** Gene hits from NCBI BLAST v2.2.25 after querying the unsexed reference genome with candidate sex determination genes on the SDR. This data has been filtered on a bit score of >100, an E-value of >1e-10 and an alignment length of >99.

| Gene identifier | Gene name | Scaffold | Identity | Align length | Mismatches | Gaps | qstart | qend | sstart | send | E-value | Bit score |
| --- | --- | --- | --- | --- | --- | --- | --- | --- | --- | --- | --- | --- |
| XM_020703654. | Cyp19b | 000374 | 78.98 | 176 | 25 | 12 | 1561 | 1730 | 1804 | 1635 | 9.00E-22 | 110 |
| XM_011475857. | Cyp19b | 000374 | 78.98 | 176 | 25 | 12 | 1567 | 1736 | 1804 | 1635 | 9.00E-22 | 110 |
| XM_020703653. | Cyp19b | 000374 | 78.98 | 176 | 25 | 12 | 1464 | 1633 | 1804 | 1635 | 8.00E-22 | 110 |
| XM_011475857. | Cyp19b | 001800 | 83.69 | 141 | 19 | 4 | 1128 | 1266 | 412 | 274 | 7.00E-28 | 130 |
| HQ449733.1 | Cyp19a1 | 001800 | 88.07 | 109 | 11 | 2 | 472 | 579 | 631 | 524 | 7.00E-28 | 128 |
| HQ449733.1 | Cyp19a1 | 001800 | 84.85 | 165 | 19 | 6 | 189 | 350 | 1494 | 1333 | 7.00E-38 | 161 |
| XM_020703653. | Cyp19b | 001800 | 83.69 | 141 | 19 | 4 | 1025 | 1163 | 412 | 274 | 6.00E-28 | 130 |
| AY273211.1 | Cyp19a1 | 001800 | 87.43 | 175 | 16 | 6 | 946 | 1117 | 412 | 241 | 6.00E-48 | 196 |
| XM_020703654. | Cyp19b | 001800 | 81.5 | 173 | 23 | 9 | 1128 | 1295 | 412 | 244 | 5.00E-29 | 134 |
| NM_001105093. | Cyp19b | 001800 | 81.93 | 166 | 22 | 8 | 951 | 1112 | 412 | 251 | 5.00E-29 | 134 |
| AY273211.1 | Cyp19a1 | 001800 | 79.87 | 149 | 24 | 6 | 400 | 545 | 1755 | 1610 | 4.00E-20 | 104 |
| AY273211.1 | Cyp19a1 | 000374 | 82.77 | 238 | 31 | 10 | 1111 | 1343 | 2359 | 2127 | 3.00E-50 | 204 |
| NM_001105093. | Cyp19b | 000374 | 79.43 | 175 | 26 | 10 | 1366 | 1535 | 1804 | 1635 | 2.00E-23 | 115 |
| AY273211.1 | Cyp19a1 | 000374 | 78.82 | 255 | 43 | 10 | 1355 | 1602 | 1810 | 1560 | 2.00E-37 | 161 |
| AY273211.1 | Cyp19a1 | 001800 | 82.8 | 157 | 21 | 6 | 249 | 402 | 2010 | 1857 | 1.00E-29 | 135 |
| AY273211.1 | Cyp19a1 | 001800 | 81.41 | 156 | 23 | 6 | 555 | 707 | 1494 | 1342 | 1.00E-25 | 122 |

### Sex marker development and validation

In total, three types of markers were designed and tested based on the *Cyp19a1a* candidate gene (n=5, Supplementary Table 2), using Y-allele (n=15, Supplementary Table 3) and specific amplification and HRM (n=15, Supplementary Table 4), respectively. All of the 15 HRM markers successfully amplified PCR products, however none of them were scorable or linked to sex based on a subset of eight samples (4 males and 4 females). Of the 15 Y-allele specific markers, one of them gave a clear PCR amplification product that was only present in the males and absent for the females. Of the 96 trevally for which the sex was confirmed by gonadal histology, all agreed with the prediction from this Y-allele specific marker (Figure 2A). Of the markers designed by amplifying large fragments of the *cyp19* candidate gene, two amplified a PCR product present in all males and absent from the females (Figure 2B).

## Discussion

In contrast to mammals and birds, cold-blooded vertebrates, and in particular teleost fishes, show a variety of strategies for sexual reproduction (Heule et al. 2014). Here we present the first near-complete genome assembly of New Zealand trevally (*Pseudocaranx georgianus*) and one of the first for a carangid species, and use this and whole-genome variation of males and females to identify sex linked regions. A genome assembly for California yellowtail (*Seriola dorsalis*) was developed, however, it was not resolved into pseudo-chromosome scale scaffolds (Purcell et al. 2018). Our assembly covers the 24 chromosomes expected from the family Carangidae and is highly-contiguous and only includes a small proportion of scaffolds that could not be anchored to any of the 24 chromosomes. The trevally genome will be useful to assemble other Carangidae fragmented genomes based on synteny, such as other *Seriola* spp. which are economically important for aquaculture around the globe (Corriero et al. 2021). We demonstrate the usefulness of this genome assembly and annotation for mapping the SD locus, which supports a XX/XY model for trevally and enabled us to develop robust PCR-based markers for sex identification in this species. We discuss these findings and outline resulting applications and implications, and provide insights how our results improve the overall understanding of the genetics of sex determination in teleost fishes.

We chose two strategies to reveal the genomic regions linked to SD in trevally. First, we screened for genomic variants that were commonly, or always, in the heterozygous state in one sex and a homozygous state in the other. We discovered two regions with high numbers of variants seen in the heterozygous state in all seven male fish assessed, which were homozygous in all six female fish assessed. One region was approximately 6 kb and located at the proximal end of a chromosome-scale scaffold (Chromosome 21), while the other region spans most of a short 3.7 kb scaffold (scaffold_000374). The second strategy was based on read-depth variation between the sexes. We found higher read-depth in males compared to females along the same two scaffolds. Because the re-sequenced females showed some deletions compared to the reference Trevally_v1 assembly, we now hypothesise that the unsexed juvenile fish used as a specimen for genome assembly must have been a male. Our results also underscore the need for studies to go beyond SNPs in their data analysis and to include the wider spectrum of structural genomic variants, including copy number repeats such as insertions, duplication and deletions, as well as fusions, fissions and translocations, to increase the power of SD detection and to better detail the full extent of sexually divergent regions (Wellenreuther et al. 2019; Mérot et al. 2020). An increasing number of studies, including on teleost species (Catanach et al. 2019), reveal that structural genomic variants encompass more genome-wide bp variants compared to SNPs, and thus hold an enormous potential to act as a potent substrate in processes involved in the eco-evolutionary divergence of species.

The region linked to sex determination on the pseudo-chromosome scaffold_001800 (Chromosome 21) is small (∼6 kb), and could have been easily missed with other methods involving less comprehensive variant detection, such as reduced-representation genotyping by sequencing. This illustrates how our strategy, using a full genome assembly coupled with the full re-sequencing of sexed individuals, efficiently enabled us to pinpoint this region, develop sex-specific markers, and identify a candidate gene. Interesting, the sex-linked short scaffold_000374 may be unanchored due to difficulties in resolving the genome assembly in the SD region. The divergence between the Y and X alleles may have prevented the Meraculous assembler from collapsing both haplotypes. Long read sequencing and a phased assembly would be useful to resolve this issue in the future.

Our results provide strong evidence that two small genomic regions form the major part of the SD locus of trevally. The presence of a *Cyp19b*-like gene within these sex-associated regions, strongly implicates a role of this gene in the sex determination of this species. No reads from female fish aligned to the gene sequence and male-specific PCR amplification of markers based on the *Cyp19b*-like gene indicates that it is specific to male fish and suggests it might play a role in the masculinisation of genetically male fish. Further research is required to elucidate what the role of *Cyp19b*-like is, and better understand its function in SD gene of trevally. Previous research has demonstrated that *Cyp19* catalyses the irreversible conversion of the androgen’s androstenedione and testosterone into the oestrogens estrone and estradiol, respectively (Piferrer and Blázquez 2005).

Recent genomic investigations have detailed that the two variants of the *Cyp19 gene* (cyp19a1a and cyp19b) were derived from the teleost specific whole genome duplication (3R) and evolved through sub-functionalization (Lin et al. 2020). Variant A (cyp19a1a) is restricted to the gonads (mainly the ovary), whereas the B variant (cyp19b) is expressed in the brain and the pituitary (Kazeto et al. 2003). When looking at studies of the genus *Seriola*, which is in the same family as trevally, variant A is only expressed in the ovaries (Koyama et al. 2019). For males, the presence of this gene appears to be related to spermatogenesis and testicular development in some species (Schulz and Miura 2002), something that is also found in other vertebrates species outside of teleosts (Robertson et al. 1999). Stage-specific gene expression during spermatogenesis in European bass (*Dicentrarchus labrax)* gonads, for example, has revealed that cyp19a1a at lower levels has a regulatory effect at the initial stages of spermatogenesis (Viñas and Piferrer 2008). In addition to this regulatory effect, cyp19a1a has also been implicated in the differentiation of sex in black porgy (*Acanthopagrus schlegeli)*, where high levels were expressed during early testicular development (Wu et al. 2008b). Females still have higher expression than males of this gene at any ontogenetic stages, however. This is probably because next to regulation and differentiation of the ovary at higher levels during early sex differentiation (Kwon et al. 2001), for females this gene is also an important factor in the female reproductive cycle (Guiguen et al. 1999).

Variant B, which resides mostly in the brain, is attributed to the control of reproduction and behaviour related to sex. RT-PCR analysis of the hermaphroditic mangrove killifish (*Rivulus marmoratus)* showed that cyp19b is expressed in both the male and hermaphroditic fish, whilst cyp19a was completely absent in males *(Lee et al. 2006)*. In addition, a study where cyp19b levels were artificially lowered in male guppy (*Poecilia reticulate)* showed these fish experience a reduction in the performance of male specific behaviours (Hallgren et al. 2006). Females also express cyp19b, but this expression is mainly restricted to the period around spawning. Work on both zebrafish (*Danio rerio*) and channel catfish (*Ictalurus punctatus*) show an increase in cyp19b right before the onset and during spawning, while a decrease and low levels cyp19b are found outside of the reproductive period (Kazeto et al. 2003). Taken together, these studies are consistent with a cyp19b being more male linked compared to cyp19a, and conversely, that cyp19a is associated with female phenotypes.

Sex chromosomes in teleosts, can either be distinguishable cytologically (heteromorphic) or appear identical (homomorphic). In both cases, one sex is heterogametic (possessing two different sex chromosomes and hence producing two types of gametes) and the other one homogametic (a genotype with two copies of the same sex chromosome, producing only one type of gamete). A male-heterogametic system is called an XX-XY system, and female-heterogametic systems are denoted as ZZ-ZW, and both types can be found side by side in closely related species (Heule et al. 2014). Close relatives of trevally show the ZW/ZZ type of sex-determination; e.g. the Japanese amberjack (*Seriola quinqueradiata*). Evidence for a ZW/ZZ type of sex-determination would come from a higher number of heterozygous SNPs in females combined with a higher number of deletions in males (the latter would be hinting at a lack of the W-chromosome) (Fuji et al. 2010). Yet in trevally the opposite is seen. When examining the SD region between the sexes, we found that in all instances males were heterozygous while all females were homozygous. A similar pattern was seen in the number of deletions. Of the 418 deletions detected, all of these were located in females, whereas none was located in males. Taken together, this all strongly indicates that trevally has XX-XY sex determining system.

Other teleost fish with a similar XX-XY sex determining mechanism have been well described. In the Atlantic cod (*Gadus morhua*), studies found deletions in females (hinting lack of a Y-chromosome) and male showed high SNP heterozygosity on the sex determination gene zKY (Kirubakaran et al. 2016) (confirmed with diagnostic PCR). In Medaka (*Oryzias latipes*) the XX-XY sex chromosomes were determined using genetic crosses and the tracking of sex linked markers (Matsuda et al. 2002). Recent studies have also revealed the putative sex gene for two carangids, the greater amberjacks (*Seriola dumerili*) and Californian yellowtails (*Seriola dorsalis*), (Fuji et al. 2010). Biochemical analyses in greater amberjacks showed a missense SNP in the Z-linked allele of 17β-hydroxysteroid dehydrogenase 1 gene (Hsd17b1) (Viñas and Piferrer 2008). In Californian yellowtails, Hsd17b1 was found in the SDR, identified by deletions in the female sex, like the SDR in trevally, however, females and not males were heterogametic in yellowtails (Purcell et al. 2018). The Hsd17b1 gene catalyzes the interconversion of estrogens (estrone<->estradiol) and androgens (androstenedione<-> and testosterone). The Hsd17b1 gene can thus be classified as an estradiol-synthesizing sex determination gene, just like cyp19 (Fuji et al. 2010), because cyp19 converts androgens to estradiol (testosterone->estradiol).

### Conclusions

As a greater number of fish genomes are sequenced, it is likely that more genes involved in the regulation of sex will discovered. This will provide much needed data for future comparative genomic work to track the evolutionary processes and patterns governing sex evolution across close and distant teleosts lineages. Given the importance of trevally and other carangid species for aquaculture production (e.g. *Seriola*) and wild fisheries (Papa et al. 2020), our reference genome will contribute to accelerating marker-assisted breeding programs, and will aid genomics-informed fisheries management programmes, by providing insights into sex ratios and sex specific effects (Bernatchez et al. 2017). This genome assembly for trevally will be a substantial resource for a variety of research applications such as population genomics and functional genomics, in both cultured and wild populations of this and other carangid species. The developed resources will future studies into teleost evolution, specifically the evolution of sex determination, which has proven to be a complex and highly variable trait in fish.

## Acknowledgements

We would like to acknowledge the PFR staff that assisted with the breeding and husbandry operation of the trevally populations; in particular Warren Fantham who oversees the larvae rearing of finfish and Therese Wells who manages the post-juvenile trevally husbandry. We would also like to thank Associate Professor P. Mark Lokman, University of Otago, for his assistance with the histological processing of gonadal tissues. This research was funded through the MBIE Endeavour Programme “Accelerated breeding for enhanced seafood production” (#C11X1603) to MW.

## Supplementary data

**Supplementary Figure 1.**
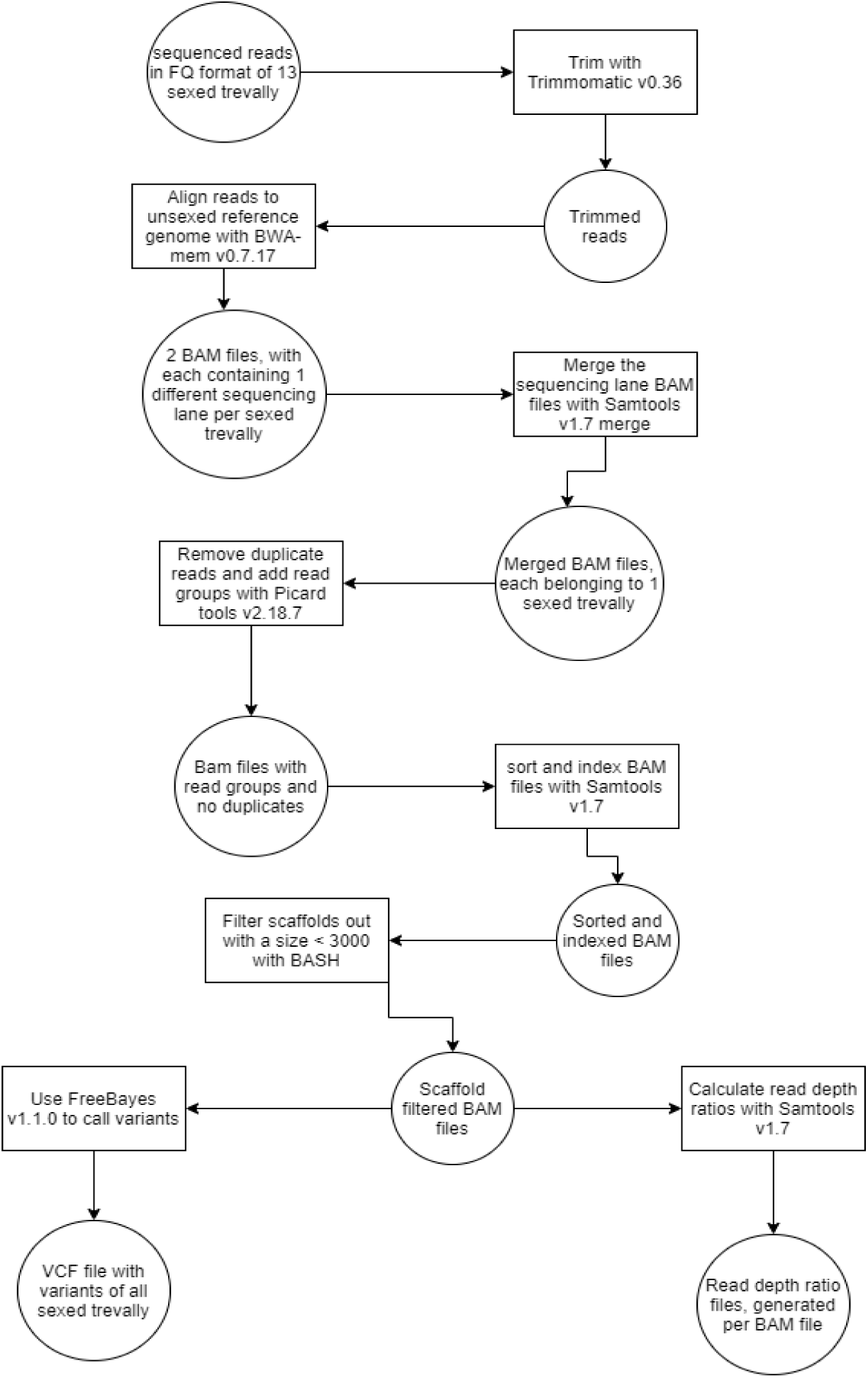
The overview of generating read depth and variant data from the sequences genomes of 13 sexed trevally.

**Supplementary Table 1.**
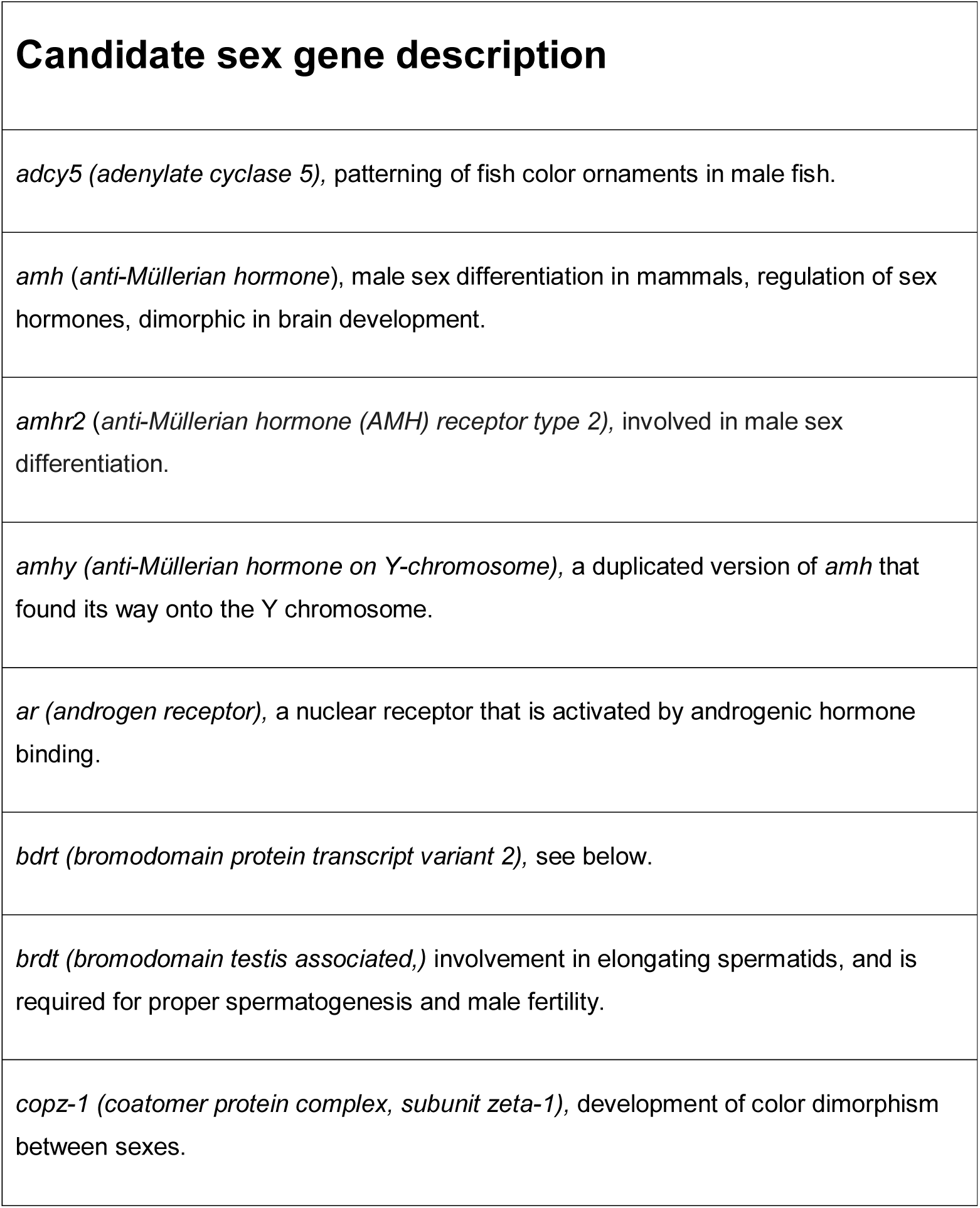
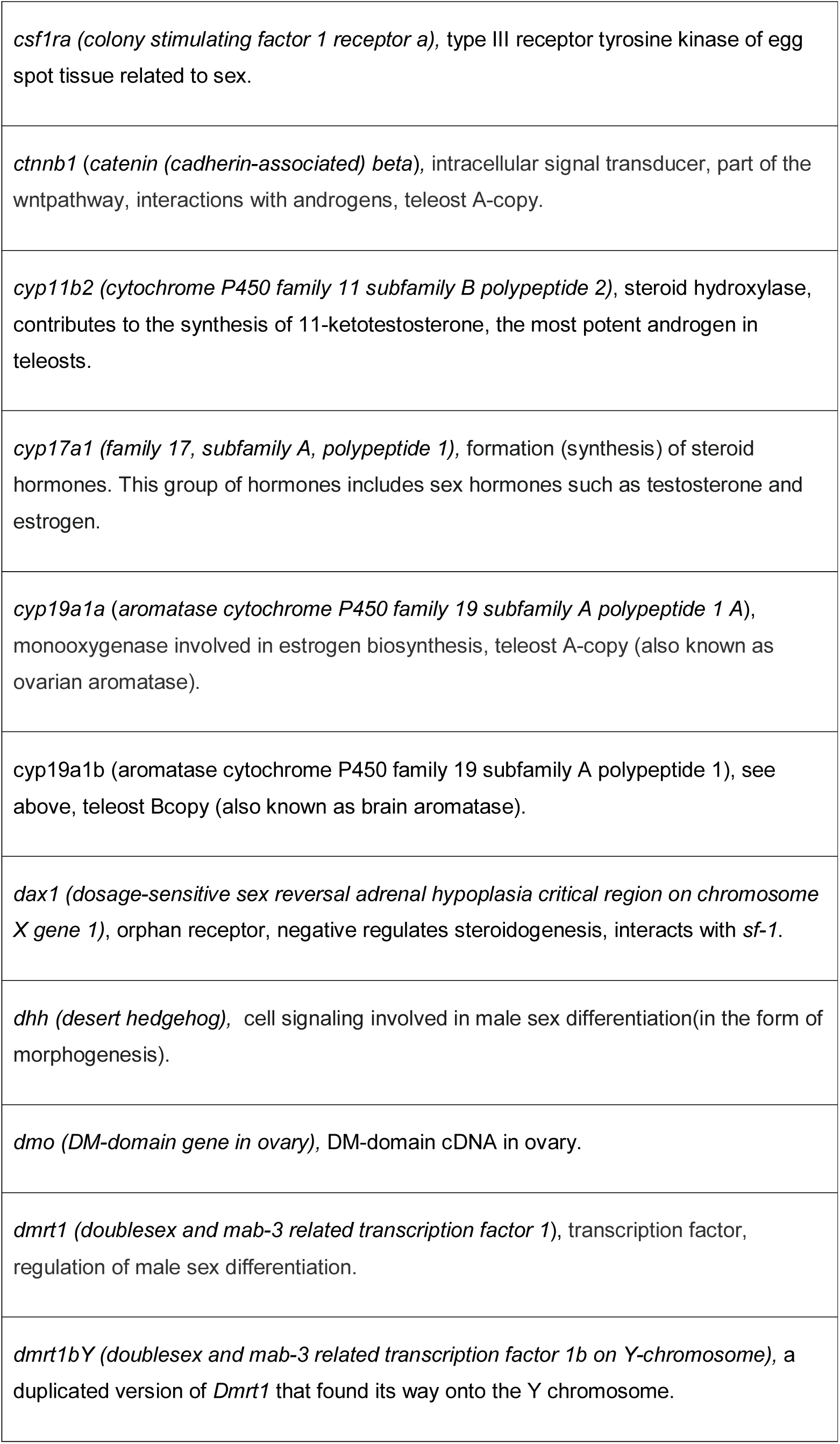
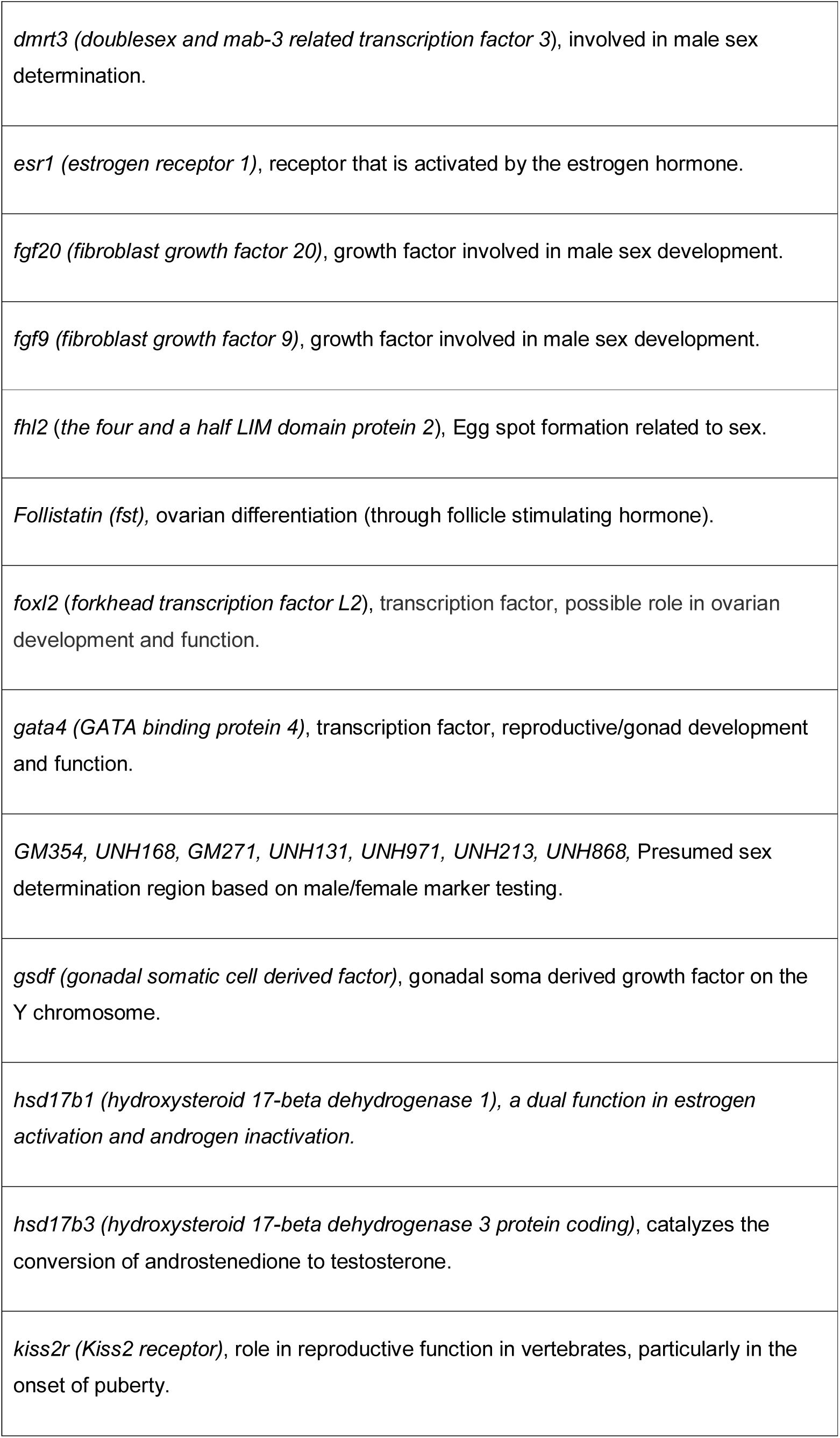
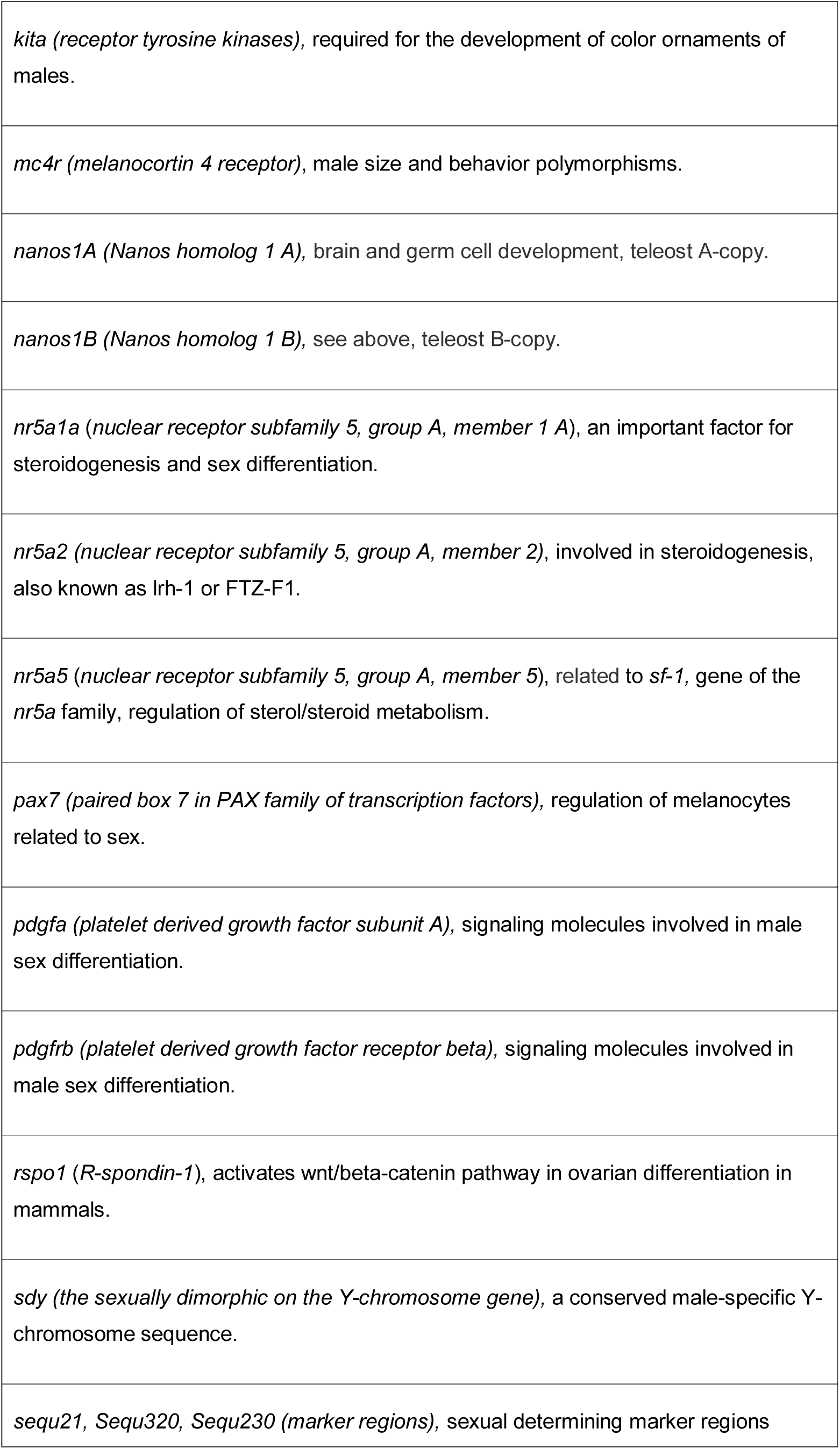
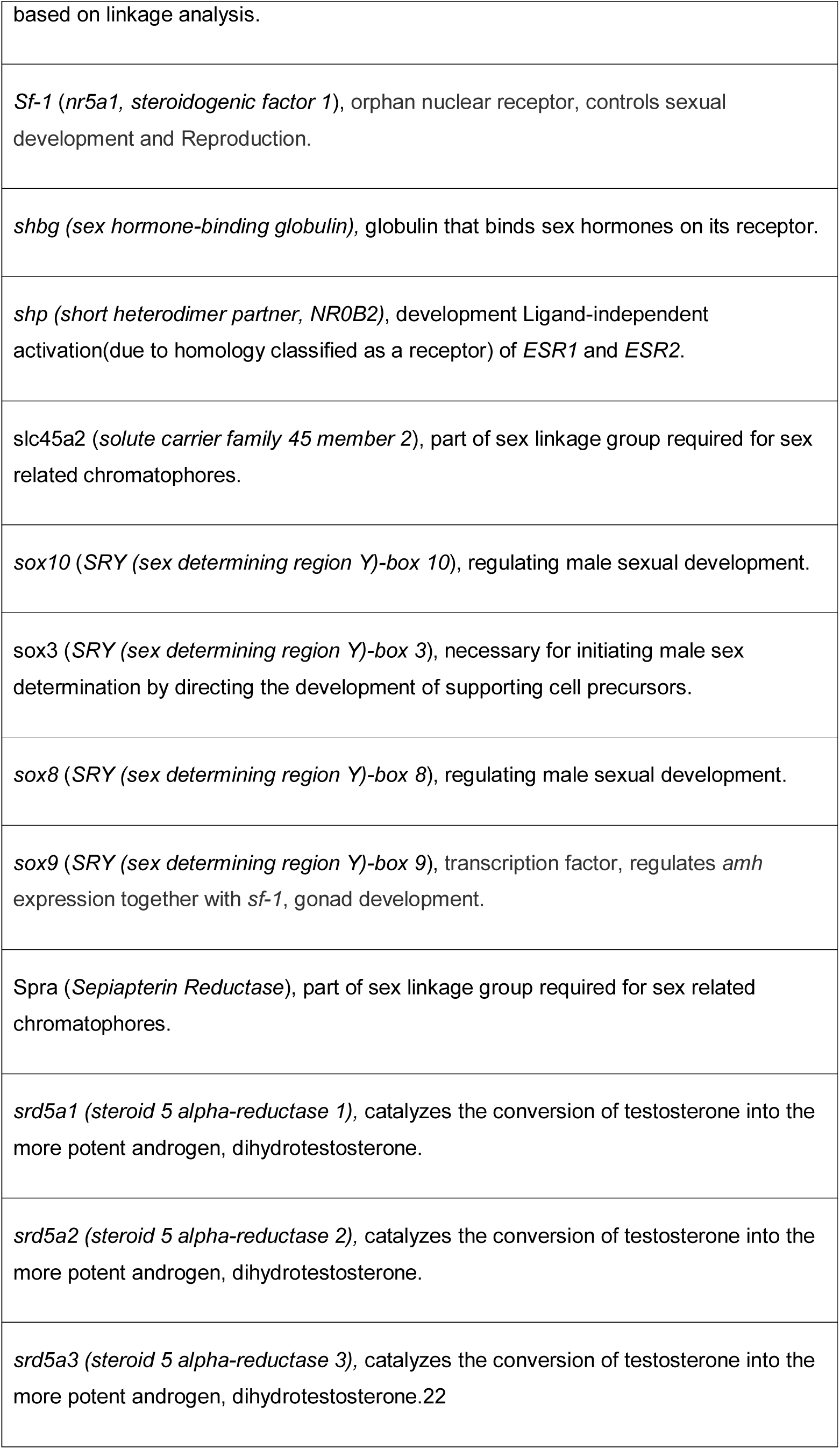
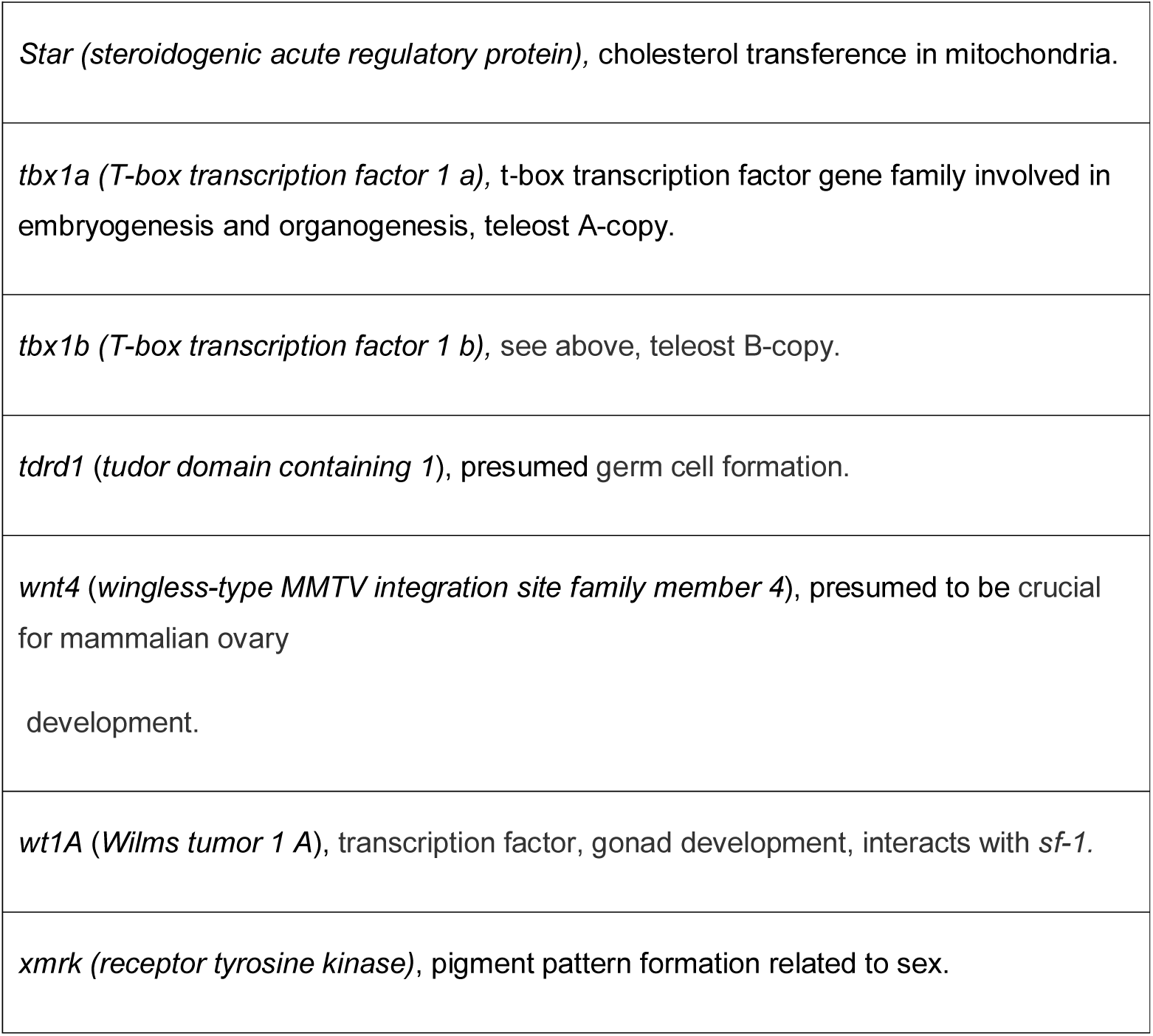
Description of the 64 sex candidate genes mined from literature (Gutbrod and Schartl 1999; Guan et al. 2000; Nanda et al. 2002; Wang et al. 2002; Yokoi et al. 2002; Lee et al. 2004; Miguel-Queralt et al. 2004; von Hofsten and Olsson 2005; Salzburger et al. 2007; Vizziano et al. 2007; Ijiri et al. 2008; Wu et al. 2008a; Roberts et al. 2009; Fuji et al. 2010; Lampert et al. 2010; Gunter et al. 2011; Yano et al. 2011; Hattori et al. 2012; Kamiya et al. 2012; Myosho et al. 2012; Nocillado et al. 2012; Yano et al. 2012; Böhne et al. 2013; Dooley et al. 2013; Forconi et al. 2013; Kottler et al. 2013; Santos et al. 2014; Úbeda-Manzanaro et al. 2016; Purcell et al. 2018). All candidate genes are sorted alphabetically on the gene sex region description column.

**Supplementary Table 2.**
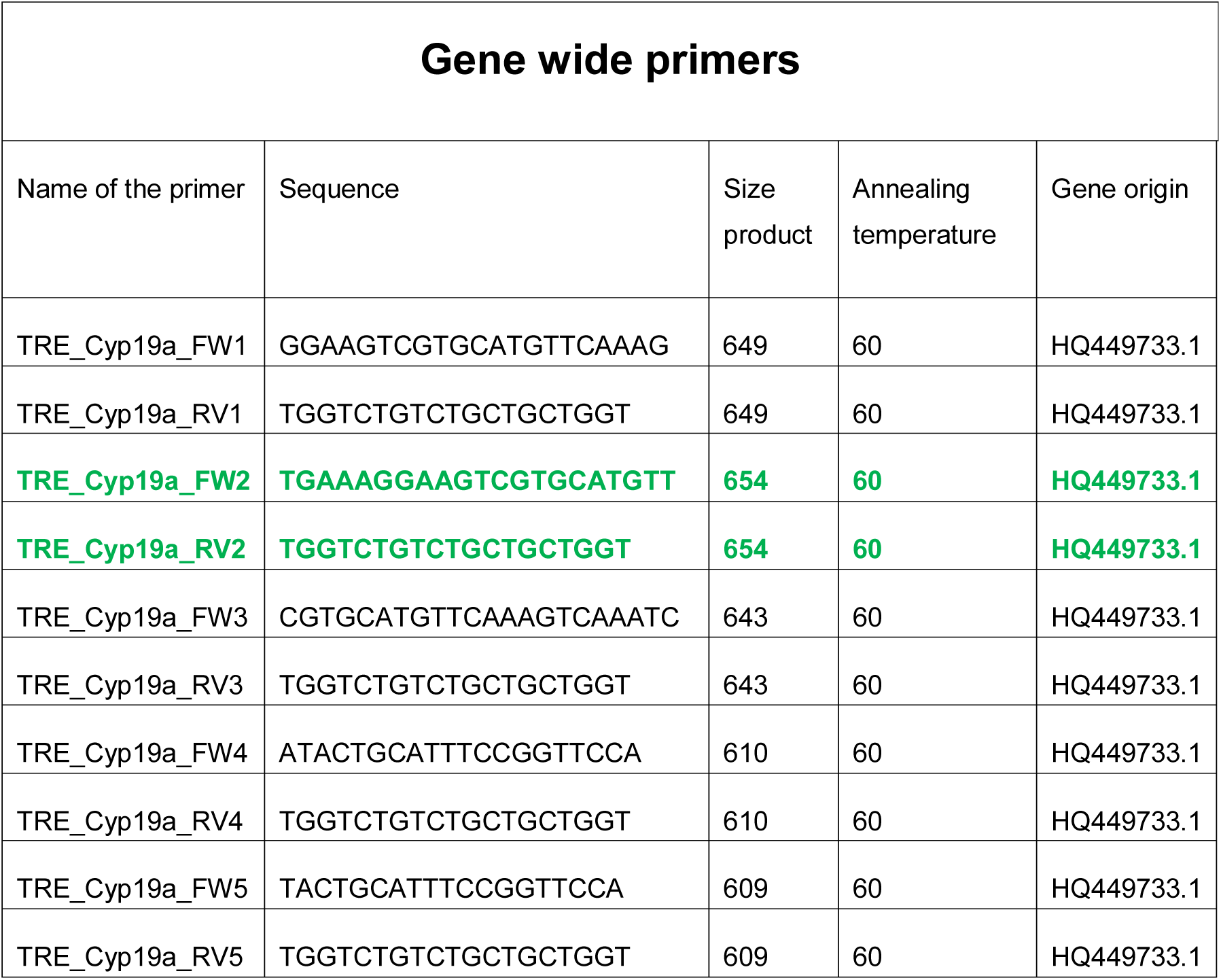
Primers names, sequence information, expected product sizes and annealing temperatures, for validation PCR. Accession numbers of orthologous genes at NCBI of the gene-wide designed primers are given. The cells with bold green letters represent the primers that were successful.

**Supplementary Table 3.**
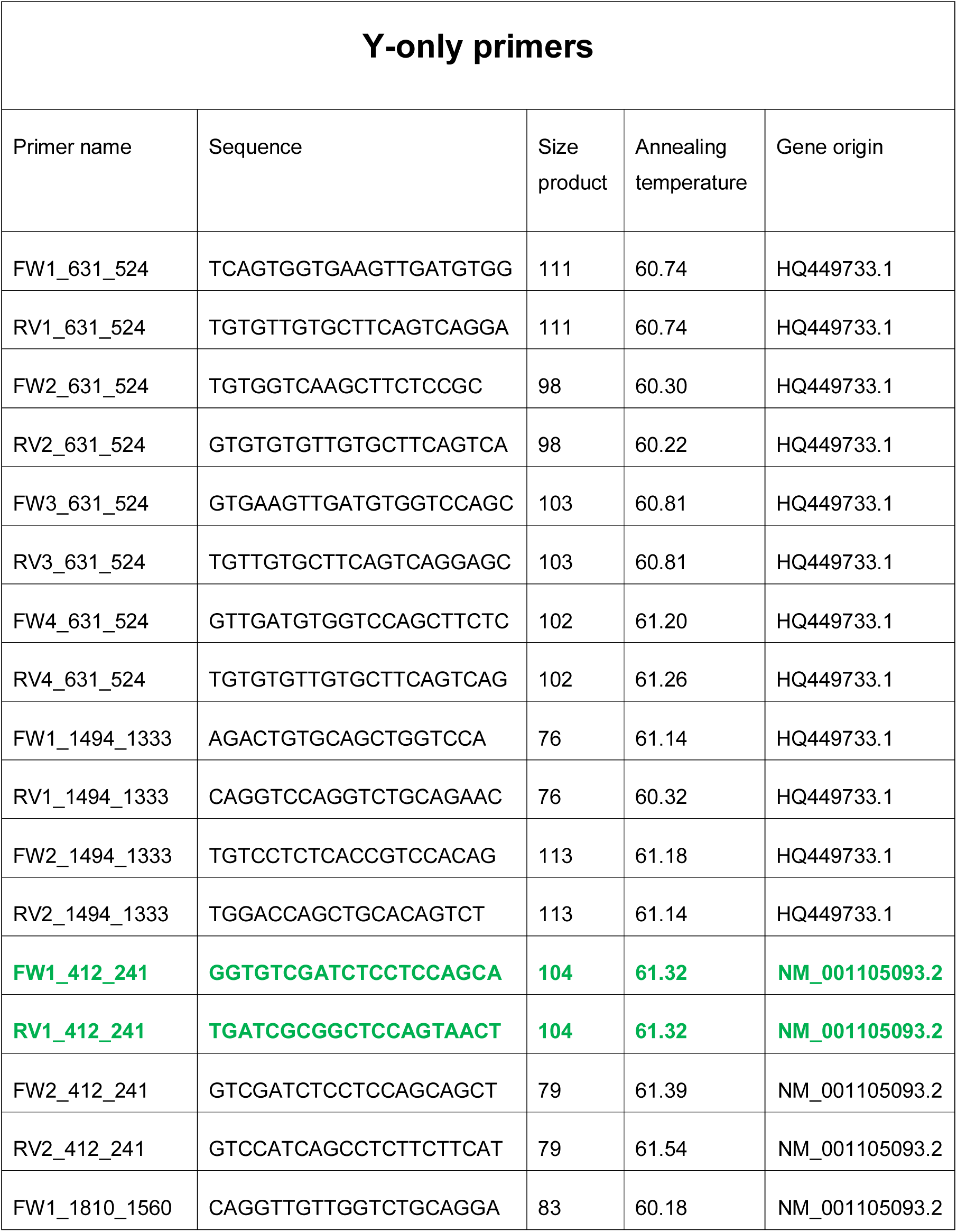
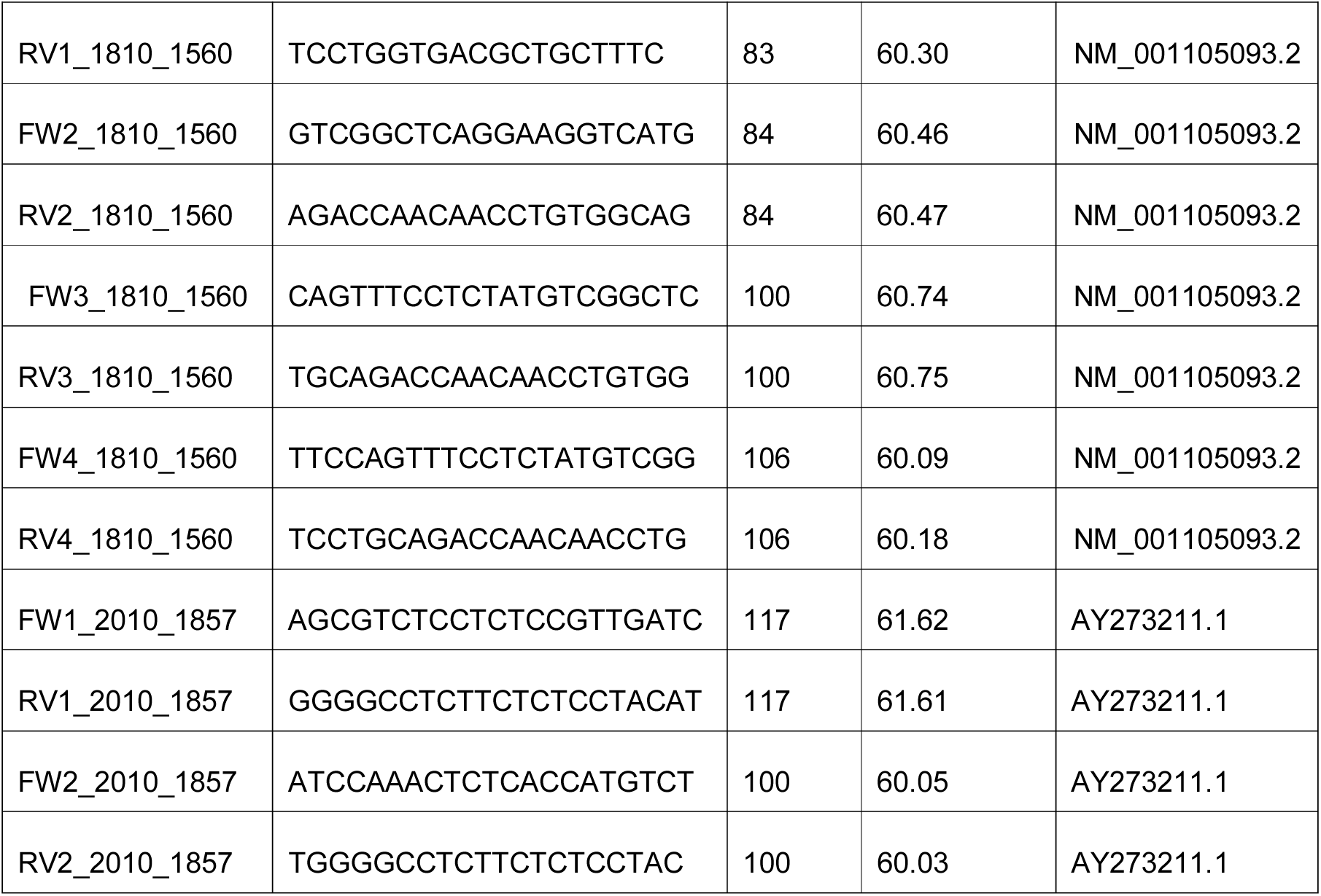
Primers names, sequence information, expected product sizes and annealing temperatures for PCR testing. The gene origin accession codes of the Y-only (Y-chromosome parts). The cells with bold green letters represent the primers that were successful.

**Supplementary Table 4.**
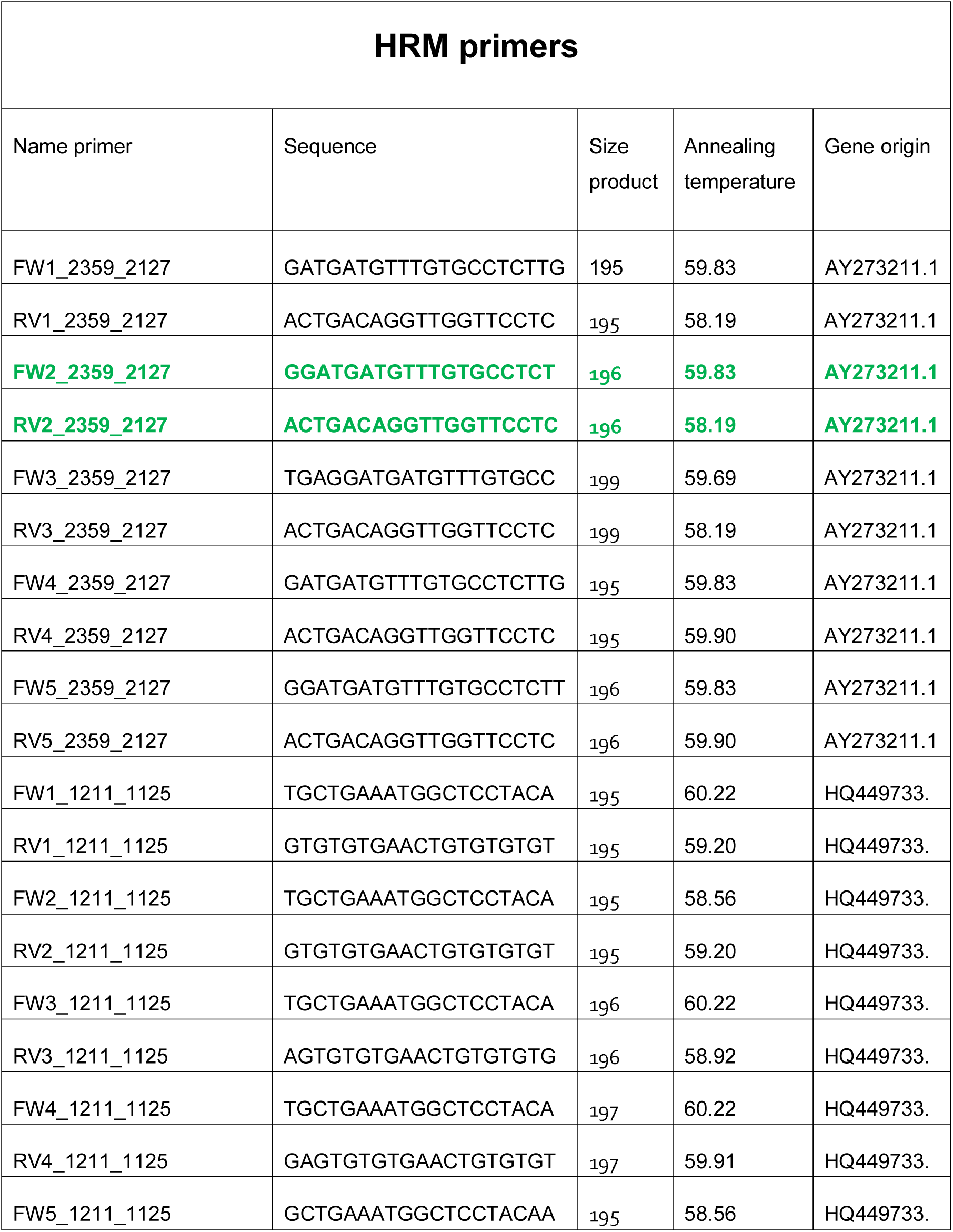
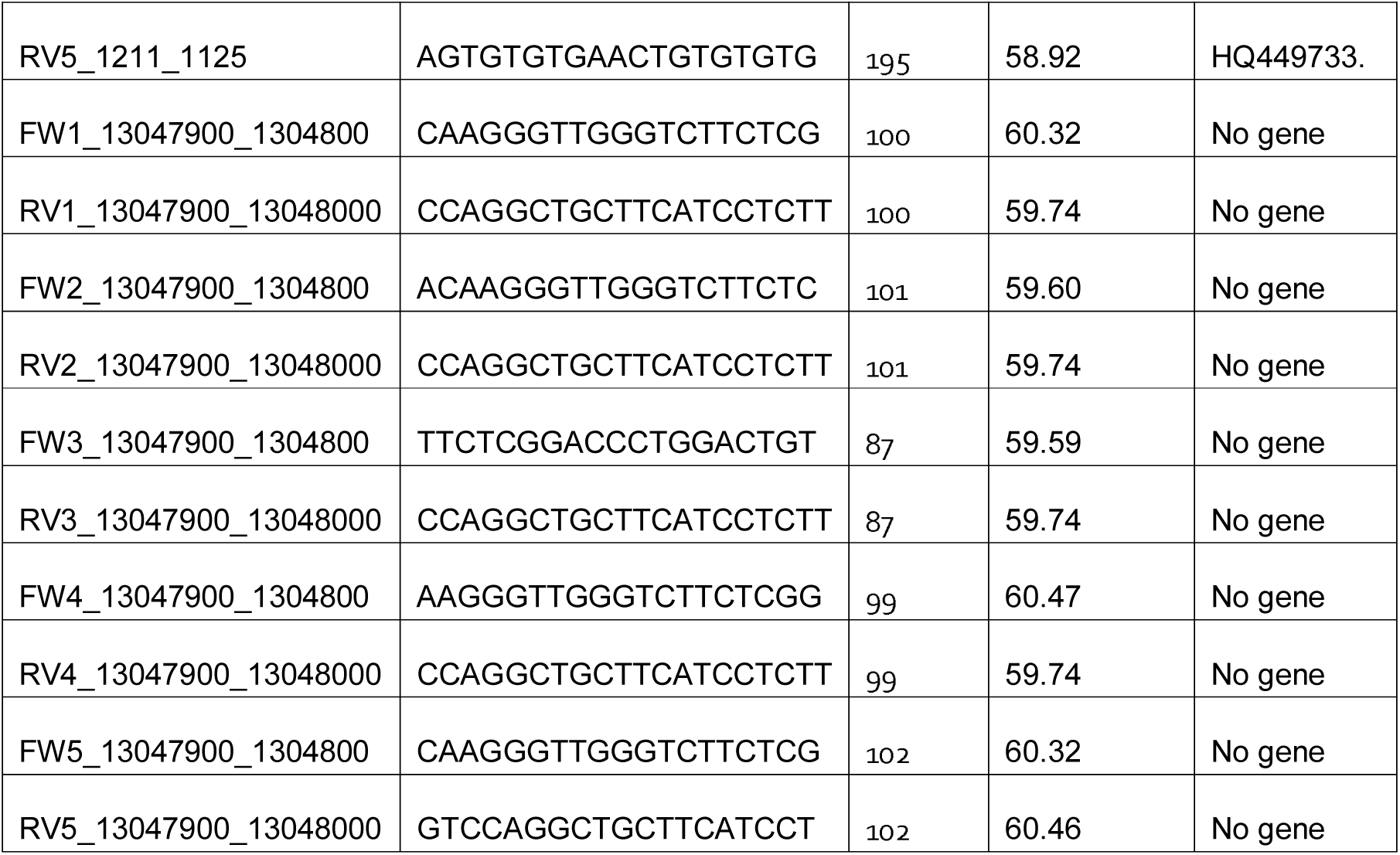
Primer names, sequences, size of product, annealing temperature for PCR and gene origin accession code of the HRM designed primers for validation. The cells with bold green letters represent the primers that turned out to be successful markers.

## Notes

https://www.genomics-aotearoa.org.nz/data

### Competing Interest Statement

The authors have declared no competing interest.

